# Mitochondrial Translation Inhibition Uncovers a Critical Metabolic-Epigenetic Interface in Renal Cell Carcinoma

**DOI:** 10.1101/2025.05.07.652786

**Authors:** Kazumi Eckenstein, Beyza Cengiz, Matthew E.K. Chang, Jessie May Cartier, Mark R. Flory, George V. Thomas

## Abstract

**Background/Objectives:** Renal cell carcinoma (RCC) exhibits distinctive metabolic vulnerabilities that may be therapeutically targeted. This study investigates how tigecycline, an FDA-approved antibiotic that inhibits mitochondrial translation, affects RCC cells and explores potential combinatorial approaches to enhance its efficacy.

**Methods:** We employed comprehensive metabolomic profiling, subcellular proteomics, and functional assays to characterize the effects of tigecycline on RCC cell lines, patient-derived organoids, and xenograft models. The synergistic potential of tigecycline with the histone deacetylase inhibitor entinostat was evaluated using combination index analysis.

**Results:** Tigecycline selectively inhibited mitochondrial translation in RCC cells, reducing mitochondrially-encoded proteins while sparing nuclear-encoded components, profoundly disrupting mitochondrial bioenergetics and reducing tumor growth in xenograft models. Subcellular proteomic analyses revealed that tigecycline treatment triggered significant accumulation of multiple histone variants concurrent with cell cycle arrest. Based on this discovery, combined treatment with tigecycline and entinostat demonstrated remarkable synergism across RCC cell lines and patient derived organoids.

**Conclusions:** Our findings identify a promising therapeutic opportunity by targeting the crosstalk between mitochondrial function and epigenetic homeostasis in RCC, with potential for rapid clinical translation given the established pharmacological profiles of both agents.

## Introduction

Metabolic reprogramming represents a compelling therapeutic avenue in oncology, as these adaptations fundamentally support neoplastic proliferation and dissemination through enzymatic activities susceptible to pharmacological intervention. Renal cell carcinoma (RCC) exhibits particularly distinctive metabolic aberrations intrinsically linked to its pathogenesis. The genetic evidence for this relationship is compelling: germline mutations in fumarate hydratase (FH), a critical tricarboxylic acid (TCA) cycle enzyme, create a predisposition specifically to renal malignancies, highlighting the unique relationship between metabolic dysregulation and kidney cancer development. Similarly, von Hippel-Lindau (VHL) tumor suppressor inactivation, observed in approximately 80% of clear cell renal cell carcinoma (ccRCC) cases, profoundly disrupts the homeostatic balance between glycolytic and oxidative phosphorylation pathways essential for cellular bioenergetics [1].

Mitochondria function as central metabolic integrators, where diverse nutrient substrates undergo complete oxidation through acetyl coenzyme A (acetyl-CoA) entry into catabolic pathways. The primary mitochondrial metabolic networks—including the TCA cycle, β-oxidation of fatty acids, electron transport chain (ETC) operation, and oxidative phosphorylation (OXPHOS)—collectively facilitate macromolecule degradation coupled to ATP synthesis. Beyond bioenergetics, mitochondria serve as biosynthetic organelles, generating precursors for cellular constituents while orchestrating nuclear gene expression in response to metabolic perturbations. The oxidative metabolism of glucose through glycolysis and the TCA cycle generates ATP while providing essential carbon skeletons for anabolic processes and maintaining cellular redox homeostasis [2,3]. Based on these integrated functions, mitochondria represent critical metabolic hubs essential for neoplastic survival and proliferation, suggesting that targeting specific mitochondrial proteins could effectively impede malignant progression while potentially overcoming resistance mechanisms to current therapies.

Tigecycline, a *glycylcycline* antibiotic, selectively inhibits mitochondrial translation by binding to the 30S ribosomal subunit, preventing aminoacyl-tRNA association with the ribosomal A site similar to its antibacterial mechanism of action [4-8]. While clinically approved for bacterial infections, tigecycline demonstrates significant antineoplastic activity across multiple cancer types, including acute myeloid leukemia (AML), hepatocellular carcinoma, colorectal cancer, and lung adenocarcinoma through inhibition of mitochondrial oxidative phosphorylation (OXPHOS) [9-13]. Notably, tigecycline has demonstrated efficacy in cancer cells with enhanced mitochondrial biogenesis and preferentially targets cancer stem cells that rely heavily on OXPHOS, and that tigecycline selectively killed leukemic stem and progenitor cells compared to their normal counterparts through inhibition of mitochondrial translation and significantly reduced basal and maximal respiratory capacity in malignant cells [14,15]. Moreover, when combined with tyrosine kinase inhibitors like imatinib, tigecycline substantially reduced chronic myeloid leukemia stem cell populations both in vitro and in xenotransplantation models by targeting mitochondrial protein synthesis and reducing oxidative metabolism.

While tigecycline’s established human pharmacokinetic profile and toxicity parameters facilitated accelerated clinical translation, the Phase I trial of tigecycline in relapsed/refractory AML patients yielded disappointing results with no objective clinical responses despite establishing the maximum tolerated dose [16]. These clinical limitations prompted us to investigate whether combinatorial approaches might enhance tigecycline’s therapeutic potential.

In this study, we demonstrate that tigecycline effectively inhibits mitochondrial translation in RCC, causing profound metabolic disruption. Next, through comprehensive proteomic and metabolomic analyses, we discovered a previously uncharacterized connection between mitochondrial dysfunction and histone homeostasis. We further show that targeting this metabolic-epigenetic interface through combination of tigecycline with the histone deacetylase inhibitor entinostat produces remarkable synergistic effects across multiple RCC models, establishing a novel therapeutic strategy with potential for rapid clinical translation.

## Material and Methods Immunoblotting

Cells were seeded into a 6-well plate and incubated overnight at 37°C, the next day they were treated with DMSO vehicle, 25 uM Tigecycline (Cayman Chemicals #15026), 2.5 uM Entinostat (MS-275, Cayman Chemicals #13284). After 72 hr unless stated otherwise, cells were washed with ice cold PBS and then lysed with cold RIPA buffer (ThermoFisher #89901) on ice using a cell scraper. PhosSTOP phosphatase inhibitor cocktail (Millipore Sigma #4906845001) and cOmplete protease inhibitor cocktail (Millipore Sigma #4693159001) were added at the time of lysis. Lysates were incubated on ice for 10 min then centrifuged at 15,000 g for 10 min at 4°C to remove insoluble material. Protein concentrations were calculated with a BCA assay (ThermoFisher #23227) generated standard curve. Equal amounts of cell lysate (15-25ug) were prepared using NuPAGE LDS sample buffer (ThermoFisher cat# NP0007) with 50mM DTT then heated at 70°C for 10 min. Proteins were resolved using the NuPAGE Novex Mini Gel system on 4% to 12% Bis-Tris Gels (ThermoFisher #NP0336BOX) and NuPAGE MES Running Buffer (ThermoFisher #NP0002) and then transferred to a 0.45um PVDF membrane (Millipore Sigma #IPFL00010). The membrane was blocked with 5% BSA in TBST for 1 hr at room temperature and then probed with primary antibodies for 16 hr at 4°C. The membrane was subsequently washed and incubated with corresponding fluorescent secondary antibodies (LI-COR #926-32211, #926-68070) for 1 hr at room temperature and washed before imaging. The fluorescent signal was captured using LI-COR (Lincoln, NE) Odyssey Imaging System. The following primary antibodies were used for western blots: NDUFA9 (ThermoFisher #459100), SDHA (Cell Signaling Technology #11998), UQCRC1 (ThermoFisher #459140), MT-CO1 (COX-1, ThermoFisher #459600), MT-CO2 (COX-2, ThermoFisher #A6404), COX4 (Cell Signaling Technology #4844), ATP5A (Abcam #ab14748), alpha-Tubulin (Cell Signaling Technology #3873), Cell Cycle Phase Determination Antibody Sample Kit (Cell Signaling Technology #17498), phospho-DRP1 (Ser616) (Cell Signaling Technology #3455), phopsho-yH2AX (S139) (Cell Signaling Technology #2577), Acetyl-Histone H3 (Lys9/Lys14) (Cell Signaling Technology #9677), HAT1 (Cell Signaling Technology #41490).

### Immunofluorescence and Lipid Droplet Labeling

Cells were seeded in glass bottom 8-chambered slides (CellVis #C8-1.5H-N) and incubated overnight at 37°C, the next day they were treated with DMSO vehicle, 25 uM Tigecycline, 2.5 uM Entinsotat (MS-275). After 24 hours unless stated otherwise, cells were washed with PBS and fixed with 4% paraformaldehyde for 10 min at room temperature. Cells were then washed twice with PBS and permeabilized in blocking buffer containing 0.3% TritonX-100 and 10% fetal bovine serum in PBS for 15 min at 37°C, followed by incubation with primary antibody for 16 hr at 4°C. Cells were then washed three times with PBS and incubated with corresponding fluorescent secondary antibodies (ThermoFisher #A11001, #A11012) for 1 hr at room temperature followed by a 10 min room temperature incubation with 1 ug/mL DAPI and two washes in PBS. For lipid droplet labeling, fixed and permeabilized cells were incubated with 5uM BODIPY 493/503 (ThermoFisher #D3922) for 30 min at room temperature and then labeled with DAPI and washed twice before imaging. Fluorescence microscopy was performed on a Zeiss Apotome3 with optical sectioning using a 63x oil objective. The following primary antibodies were used for immunofluorescence: Citrate Synthase (Santa Cruz Biotechnology #sc-390693), phopho-yH2Ax (S139) (Abcam #ab26350).

### Oxygen Consumption Rate Assay

Oxygen consumption rate was measured using a SeaHorse XFe96 analyzer (Agilent Technologies). Cells were seeded in a SeaHorse XF96 microplate and treated with the indicated compounds for 24 hours at 37°C, after which the culture media was removed and changed to Mito Stress Test Assay Medium, pH 7.4, consisting of SeaHorse XF DMEM (Agilent # 103575-100), 1 mM pyruvate (Agilent #103578-100), 2 mM glutamine (Agilent #103579-100), and 10 mM glucose (Agilent #103577-100). The SeaHorse XF96 cartridge was then loaded with Oligomycin (Millipore Sigma #75351), FCCP (Cayman Chemicals #15218), and Rotenone (Millipore Sigma #R8875) + Antimycin-A (Millipore Sigma #A8674). Calibration and measurements were carried out according to the manufacturer’s instruction. Oxygen consumption rates were normalized to cell viability.

### Cell Culture

The human renal cell carcinoma cell lines ACHN, CAKI-1, and 786-O used in this study were obtained from the ATCC (#CRL-1611, HTB-46, CRL-1932 respectively). They were maintained in DMEM, McCoy’s 5A, and RPMI (respectively) + 10% fetal bovine serum plus 1% pen-strep at 37°C in a 5% CO2 incubator.

### Organoid Sourcing and Culture

The organoids (743489-274-T-V1-organoid, referred to as NCI-274, and 487391-300-R-V3, referred to as NCI-300) used in this study were developed by NCI PDMR. https://pdmr.cancer.gov/. Organoids were embedded in ice cold Cultrex Reduced Growth Factor Basement Membrane Extract (BME), Type 2 (R&D Systems, cat# 3533-010-02) and plated in warmed 24-well plates, then incubated for 15 minutes at 37°C to allow for BME polymerization. After polymerization, NCI media 6A or 6E was added to NCI-274 or NCI-300, respectively, as specified by PDMR SOP30101. NCI media 6A consisted of Organoid Basic Media and L-WRN conditioned media at a 1:1 ratio and supplemented with 1.25 mM N-acetylcysteine (Sigma, cat# A9165), 10 mM Nicotinamide (Sigma, cat# 72340), B-27 supplement (Thermo Fisher, cat# 17504044), N-2 supplement (Thermo Fisher, cat# 175002001), and 10 uM Y-27632 (Cayman Chemicals, cat# 10005583); NCI media 6E consisted of the components described in NCI 6A media recipe plus 50 ng/mL hEGF (R&D Systems #AFL236), 10 ng/mL hFGF-10 (R&D Systems #345-FG), 1 ng/mL hFGF-2 (R&D Systems #3718-FB), and 1 uM PGE2 (Tocris #2296). Organoid Basic Media consisted of Advanced DMEM-F12 (Thermo Fisher, cat# 12634028), 100 ug/mL Primocin (Invivogen, cat# ant-pm-05), Glutamax (Thermo Fisher, cat# 35050061), and 10 mM HEPES (Thermo Fisher, cat# 12630080); L-WRN conditioned media was made using the L-WRN cell line (ATCC, cat# CRL-3276) as specified by the manufacturer’s protocol and recommended reagents. Organoid density and size were monitored and passaged when criteria were met according to PDMR SOP40103. Organoid-BME domes were digested using 750 ug/mL Dispase II (Thermo Fisher, cat# 17105041).

### Cell and Organoid Viability Assays

Cell viability was measured using crystal violet staining assay. Cells were plated in a 96 well plate and 16 hr later, treated with indicated compound. After 72 hr unless stated otherwise, media was removed, and cells were washed once with PBS. Cells were then fixed with 4% paraformaldehyde for 10 min at room temperature, followed by two washes in PBS and incubation with 0.05% crystal violet (ThermoFisher #405831000) plus 2.5% MeOH for 30 min at room temperature. Cells were then washed twice with ddH2O and air dried for 16 hr. Cell-bound crystal violet was destained with 100% MeOH for 20 min at room temperature and absorbance at 590 nm was measured using a Cytation5 Multimode Reader (BioTek, Agilent Technologies).

Organoid viability was measured using the CellTiter-Glo 3D Cell Viability Assay (Promega #G9682). Organoids were released from BME domes before proceeding with the assay, which was performed based on the manufacturer’s instructions.

### Whole cell and mitochondrial isolation prep for Proteomics

Cells were plated in a 10 cm TC dish and 16 hr lated treated with 25 uM Tigecycline or matched DMSO vehicle control. After 72 hr, media was removed and washed once with ice cold PBS. For whole cell isolation, cells were lysed on ice with cold RIPA plus PhosSTOP phosphatase and cOmplete protease inhibitor cocktails using a cell scraper. Lysates were incubated on ice for 10 min then centrifuged at 15,000 g for 10 min at 4°C to remove insoluble material. Supernatant was flash frozen using liquid NO2 and stored at -80°C until shipment.

Mitochondria were isolated using Mitochondria Isolation Kit for Cultured Cells (ThermoFisher #89874), using a dounce homogenizer according to the manufacturer’s instructions. Mitochondria fractions were flash frozen using liquid NO2 and stored at -80°C until shipment.

### Metabolomics

Cells were plated in a 10 cm TC dish and 16 hr later treated with 25 uM Tigecycline or matched DMSO vehicle control. After 72 hr, cells were gently rinsed with freshly prepared 50 mM ammonium formate (Fisher Scientific, #A666-500), pH 6.8 buffer. A small volume of 50 mM ammonium formate, pH 6.8 was added to the dish and cells were collected using a cell scraper. Cells were transferred to a tube and centrifuged for 5 min at 1000 rpm using a traditional tabletop microcentrifuge. Remaining buffer was removed, and cell pellets were flash frozen using liquid NO2 and stored at -80°C until shipment.

Metabolites were extracted from cell pellets using MeOH:H2O 80:20 Protein Precipitation Protocol. All samples were adjusted to 200 ug total protein amount per sample and normalized by protein amount using BCA. Sample acquisition was performed using reverse phase liquid chromatography (RPLC) with a Thermo Scientific Q Exactive HF (LC-Hybrid Quadrupole-Orbitrap MS/MS) instrument in positive ion mode. Hydrophobic metabolite analysis was performed using RPLC separation over a 30 min gradient, using Hypersil Gold C18 column, Mobile phase A: 100 H2O with 0.1% FA, and Mobile phase B: 80:20 CAN:H2O with 0.1% FA. Data alignment and biostatistical analysis was made with Progenesis QI 2.0 and MetaboAnalyst 4.0.

### Mass spectrometry

We performed Data Independent Analysis (DIA) performed using an Eclipse Tribrid Orbitrap mass spectrometer (Thermo Scientific, San Jose, CA)[<u>29,30</u>]. A chromatographic library was generated using the Spectronaut integrated database search engine Pulsar to generate a hybrid using DIA and DDA spectra. The library was generated using a combination of six gas phase fractions (GPF) and full scan data-Dependent analysis of the biological sample pool. The GPF acquisition used 4 m/z precursor isolation windows in a staggered pattern (GPF1 398.4–502.5 m/z, GPF2 498.5– 602.5 m/z, GPF3 598.5–702.6 m/z, GPF4 698.6–802.6 m/z, GPF5 798.6–902.7 m/z, GPF6 898.7–1002.7 m/z) at a resolution of 60,000, AGC target was set to custom with a normalized target of 1000%, maximum injection time was set to dynamic with a minimum of nine points across the peak, and an NCE of 33 using higher-energy collision dissociation (HCD). Three DDA full scan runs were performed on the biological pool. The MS1 precursor selection range is from 375–1500 m/z at a resolution of 120K with a normalized automatic gain control (AGC) target of 250% and an automatic maximum injection time. Quadrupole isolation of 0.7 Th for MS^2^ isolation and CID fragmentation in the linear ion trap with a collision energy of 35% and a 10ms activation time. The MS^2^ AGC was in standard mode with a 35 ms maximum injection time. The instrument was operated in a data-dependent mode with a 3 second cycle time and the most intense precursor priority and the dynamic exclusion set to an exclusion duration of 60s with a 10ppm tolerance. Biological samples were run on an identical gradient as the GPFs using a staggered window scheme of 8 m/z over a mass range of 385–1015 m/z. Precursor isolation was performed in the Orbitrap at 60,000 resolution with a dynamic maximum injection allowing for a minimum of nine points across the peak, a custom AGC normalized to 1000% and an NCE of 33 using HCD. The species specific FASTA database for Mus musculus (UP000000589_10090) containing 22,282 proteins and the known contaminants (Crap_uniprot_No_human_MRSonbeadV2) were downloaded from Uniprot. Variable modifications considered were: Carbamidomethylation C. The digestion enzyme was Trypsin with a maximum of 1 missed cleavage site. Peptides with charges from 2–3 and length of 6–30 amino acids and less than or equal to an FDR of 0.01 were considered. Protein groups were thresholded to achieve a protein FDR less than 1.0%.

Protein groups and their unscaled quantified intensities generated by Spectronaut were exported and further analyzed in the statistical software R v4.3.1. Decoys, protein groups with zero quantification, and known contaminants were removed from further analyses. Standard quality control procedures were conducted including extreme outlier identification, total intensity per sample distribution, sample correlation, and principal component analysis (PCA). Then, we transformed the data using the Cyclic Loess normalization from the Linear Models for Microarray Data (limma v3.56.2) Bioconductor package. Prior to performing statistical analysis, PCA and correlation matrices (not shown) were plotted to ensure data quality. Results were adjusted for multiple testing error to calculate the q-value. Proteins were statistically significantly different with an FDR adjusted q-value < 0.05. Log2 fold-changes were also recorded but were not used to define significance. To gain further insights and identify key pathways, the results of the differential analysis were investigated with Advaita Bio’s iPathwayGuide (iPG, [17]) This web-based tool identifies significant pathways via the proprietary “Impact Analysis” method and determines enriched gene ontologies and KEGG pathways. The p-value of each KEGG pathway was then corrected for multiple comparisons using false discovery rate (FDR).

## Results

### Tigecycline Inhibits Mitochondrial Translation and Impairs Cellular Bioenergetics in RCC

First, we demonstrated that tigecycline specifically inhibited translation of mitochondrially encoded proteins cytochrome c oxidase I (MT-CO1) and MT-CO2, without affecting nuclear-encoded mitochondrial proteins COX-4, ATP5A and ubiquinol–cytochrome c reductase core protein II (UQCRC2) **(Figure 1A)**. This selective inhibition confirmed tigecycline’s on-target activity against mitochondrial protein synthesis machinery in renal cell carcinoma (RCC) cells.

**Figure 1.**
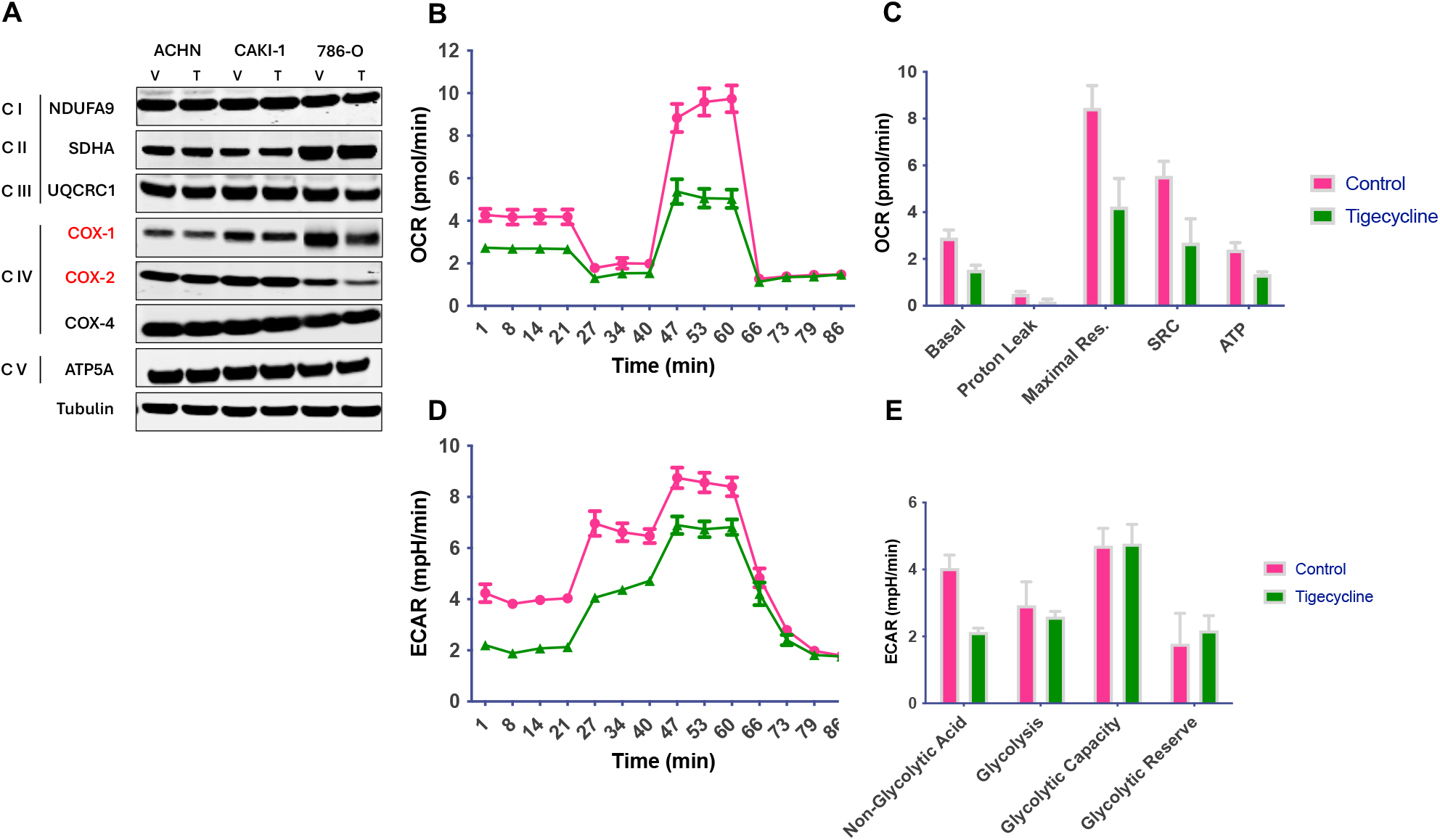
Tigecycline selectively inhibits mitochondrial translation and impairs cellular bioenergetics in RCC cells. **(A)** Western blot analysis of mitochondrially-encoded proteins (MT-CO1, MT-CO2) and nuclear-encoded mitochondrial proteins (ATP5A, UQCRC2) in ACHN, 786-O, and CAKI-1 RCC cells following treatment with tigecycline (25 μM) or vehicle control (DMSO) for 48 hours. **(B)** Representative oxygen consumption rate (OCR) traces measured by Seahorse XF analyzer in ACHN cells treated with tigecycline (25 μM) or DMSO for 24 hours. **(C)** Quantification of key bioenergetic parameters from Seahorse analysis, including basal respiration, maximal respiration, spare respiratory capacity, and ATP production. Data represent mean ± SEM from three independent experiments. **(D)** Extracellular acidification rate (ECAR) measured in RCC cells following tigecycline or DMSO treatment, showing decreased glycolytic activity in tigecycline-treated cells. **(E)** Quantification of key bioenergetic parameters from Seahorse analysis, including non-glycolytic acidification, glycolysis, glycolytic capacity and glycolytic reserve.

Next, to characterize the metabolic consequences of mitochondrial translation inhibition, we conducted comprehensive bioenergetic profiling using the Seahorse XF Analyzer [18]. Treatment with tigecycline *significantly* reduced multiple parameters of mitochondrial respiration in RCC cells as evidenced by significant reductions in basal oxygen consumption rate (OCR) and spare respiratory capacity (SRC). The diminished SRC is particularly important as it represents the cells bioenergetic reserve for responding to increased energy demands under stress conditions **(Figure 1B, C)**. ATP production was markedly decreased following tigecycline treatment, creating an energetic deficit that compromises multiple cellular processes. Unexpectedly, tigecycline also reduced extracellular acidification rate (ECAR), indicating concurrent impairment of glycolysis **(Figure 1D, E)**. This dual inhibition of major energy-producing pathways—OXPHOS and glycolysis—suggests that tigecycline disrupts metabolic compensation mechanisms that typically allow cancer cells to adapt to energetic stress.

The inability to maintain either oxidative or glycolytic ATP production likely contributes significantly to tigecycline’s antiproliferative effects in RCC cells. Having established tigecycline’s primary effects on bioenergetics, we next sought to characterize its broader impact on cellular metabolism through comprehensive metabolomic profiling.

### Tigecycline Treatment Induces Global Metabolic Reprogramming in RCC Cells

Metabolomic analysis revealed profound alterations in cellular metabolism following tigecycline treatment of RCC cells **(Figure 2A)**. Specifically, we observed a marked increase in glutathione and glutamine. Notably, the significant increase in glutathione represents a critical cellular adaptive response to oxidative stress generated by impaired electron transport chain function. Studies have shown that inhibition of mitochondrial translation by tigecycline induces reactive oxygen species accumulation [12], necessitating enhanced antioxidant defense mechanisms. The observed elevation of L-glutamine further supports this adaptation, as it serves as a precursor for glutathione synthesis and as an alternative anaplerotic substrate for the TCA cycle when mitochondrial function is compromised. We also observed a downregulation of cystine. As the oxidized dimer of cysteine, cystine is typically imported into cells via the xCT antiporter in exchange for glutamate and subsequently reduced to cysteine for glutathione synthesis.

**Figure 2.**
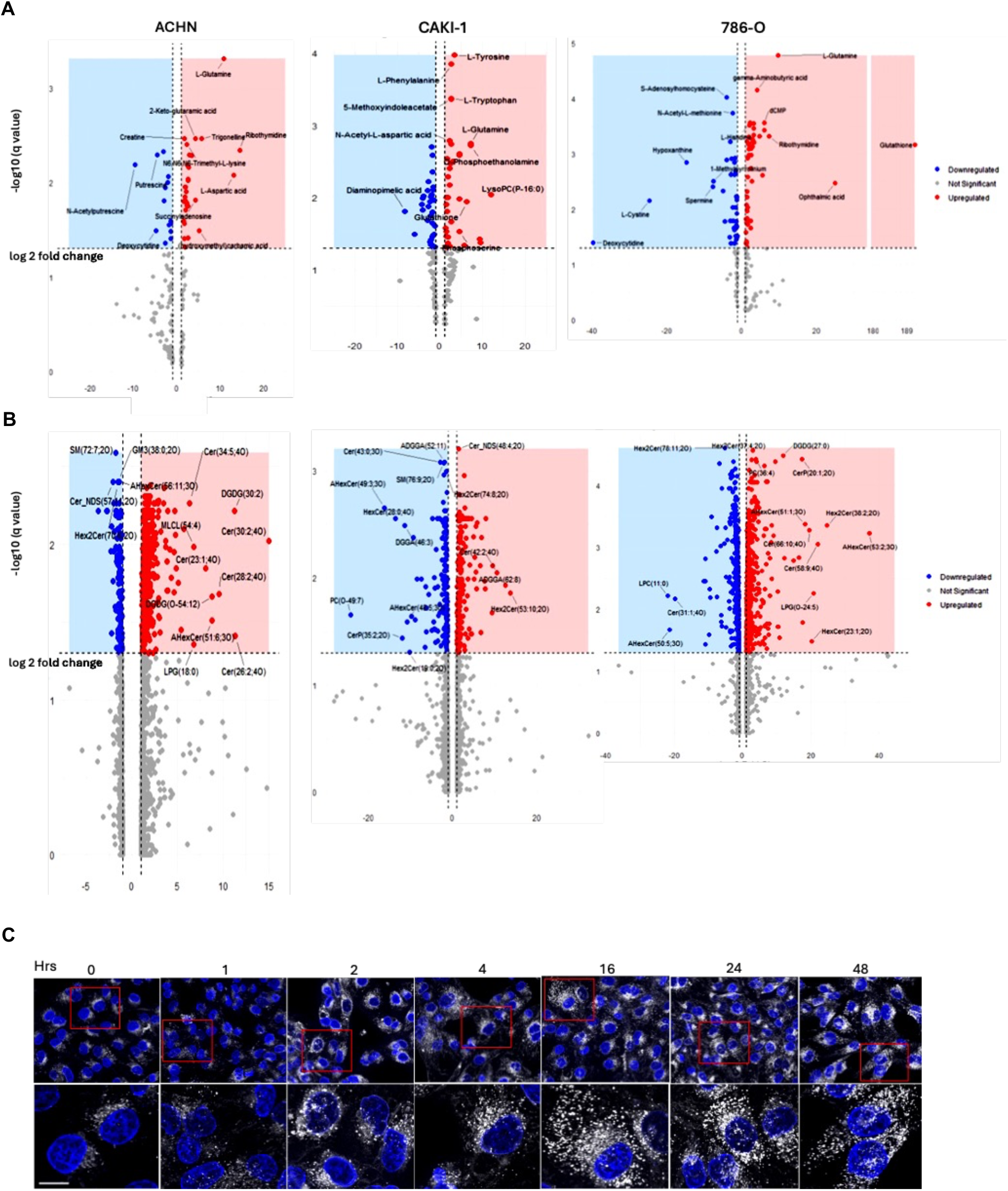
Tigecycline induces global metabolic reprogramming and lipid accumulation in human RCC cells. **(A)** Volcano plots of metabolite changes in ACHN, CAKI-1, and 786-O cells following treatment with tigecycline (25 μM) for 48 hours compared to DMSO control. Blue dots represent significantly downregulated metabolites, red dots indicate significantly upregulated metabolites (p<0.05, |log2 fold change|>1), and gray dots represent non-significant changes. Key metabolites are labeled. **(B)** Volcano plots showing lipidomic profiles of tigecycline-treated versus control RCC cells. **(C)** Time course of lipid droplet formation in ACHN cells treated with tigecycline (25 μM) for the indicated time points (0-48 hours). Upper panels show merged images of BODIPY 493/503 staining (white channel) for neutral lipids and DAPI nuclear counterstain (blue). Higher magnification of red boxed areas is shown in the lower panel for improved visualization of lipid droplet accumulation. Images were acquired using Zeiss Apotome microscopy with consistent acquisition parameters. Scale bar = 20 μm.

The decreased cystine levels, coupled with increased glutathione, indicate that cells are prioritizing the maintenance of glutathione pools through alternative pathways, possibly through enhanced transsulfuration from methionine or increased recycling of existing glutathione.

Additionally, the downregulation of hypoxanthine, putresine and spermine downregulation represents disruption of polyamine metabolism, which is critical for nucleic acid stabilization, cell proliferation, and protection against oxidative damage.

Together, these metabolic shifts reflect comprehensive reprogramming of cellular metabolism to cope with mitochondrial dysfunction, balancing between maintaining critical defense mechanisms (glutathione) while downregulating energy-intensive pathways (polyamine synthesis) under conditions of bioenergetic stress. The extensive metabolic alterations observed prompted us to investigate whether these changes were also reflected in the lipid composition of tigecycline-treated cells, as mitochondria play critical roles in lipid metabolism.

### Tigecycline Profoundly Alters Lipid Metabolism in RCC Cells

Lipidomic analysis revealed distinct patterns of lipid alteration across RCC cell lines following tigecycline treatment **(Figure 2B)**. We observed significant upregulation of ceramides (Cer), hexosylceramides (HexCer), and diacylglycerols (DGDG) was observed, with certain ceramide species showing log2 fold changes exceeding 20 (Cer(20:1-20), Cer(58:9-40)). The pronounced ceramide enrichment is particularly significant, as ceramides function as key mediators of cellular stress responses, capable of inducing cell cycle arrest and apoptosis. This lipid accumulation was further confirmed by BODIPY staining, which showed increased cytoplasmic lipid droplet formation in tigecycline-treated cells **(Figure 2C)**. Mechanistically, the increased lipid droplets may be the result of the upregulation of diacylglycerols, which serve as direct precursors for triacylglycerol synthesis, which are the primary component of lipid droplets. The impairment of mitochondrial β-oxidation following tigecycline-induced translation inhibition redirects fatty acid substrates toward storage in lipid droplets rather than oxidative metabolism. Concurrently, the pronounced ceramide accumulation contributes to lipid droplet biogenesis through activation of lipid droplet-associated proteins and further inhibition of mitochondrial fatty acid oxidation. This coordinated lipid remodeling represents a metabolic adaptation to mitochondrial dysfunction, allowing sequestration of potentially toxic lipid intermediates that accumulate when mitochondrial catabolism is compromised.

These findings indicate that tigecycline-induced mitochondrial translation inhibition drives comprehensive reprogramming of lipid metabolism, promoting accumulation of stress-responsive signaling lipids while compromising membrane structural integrity. The consistent pattern of ceramide accumulation across different RCC cell lines suggests this represents a conserved response to mitochondrial dysfunction rather than a cell line-specific phenomenon, potentially contributing to the observed cell cycle arrest and cytotoxicity through established ceramide-mediated stress signaling pathways.

### Tigecycline Modulates Mitochondrial Dynamics and Integrated Stress Response

As we observed these substantial changes in metabolism and membrane composition, we hypothesized that tigecycline might also affect mitochondrial dynamics and morphology, which are closely linked to mitochondrial function. Immunoblot analysis revealed that tigecycline treatment progressively reduced phosphorylation of dynamin-related protein 1 (DRP1) at serine 616 (p-DRP1 Ser616) in all three RCC cell lines examined **(Figure 3A)**. DRP1 is a critical GTPase that regulates mitochondrial fission, with its activity primarily controlled through post-translational modifications, particularly phosphorylation [19]. Phosphorylation at serine 616 promotes DRP1 activation and subsequent mitochondrial fragmentation, a process frequently upregulated in cancer cells to support their metabolic demands and proliferative capacity. The observed reduction in pDRP1 Ser616 following tigecycline treatment indicates significant impairment of mitochondrial fission processes. This was confirmed by the immunofluorescence images which revealed dramatic alterations in mitochondrial morphology following tigecycline treatment **(Figure 3B)**. In vehicle-treated cells (left panel), mitochondria exhibit the typical tubular, interconnected network structure with numerous branch points, appearing as punctate and reticular structures (white) distributed throughout the cytoplasm, which represents normal mitochondrial dynamics with balanced fission and fusion processes. In contrast, tigecycline-treated cells (right panel, white with red arrowheads) display extensively elongated, hyperfused mitochondrial networks with significantly reduced fragmentation. This phenotype is consistent with impaired mitochondrial fission. Concurrently, tigecycline significantly increased phosphorylation of eukaryotic initiation factor 2α (p-eIF2α Ser 51) and upregulated activating transcription factor 4 (ATF4), hallmarks of integrated stress response activation [20] **(Figure 3C)**. This stress response triggers attenuation of global protein synthesis while selectively upregulating stress-responsive proteins. The activation of these adaptive pathways reveals mechanisms by which cancer cells attempt to mitigate metabolic perturbations.

**Figure 3.**
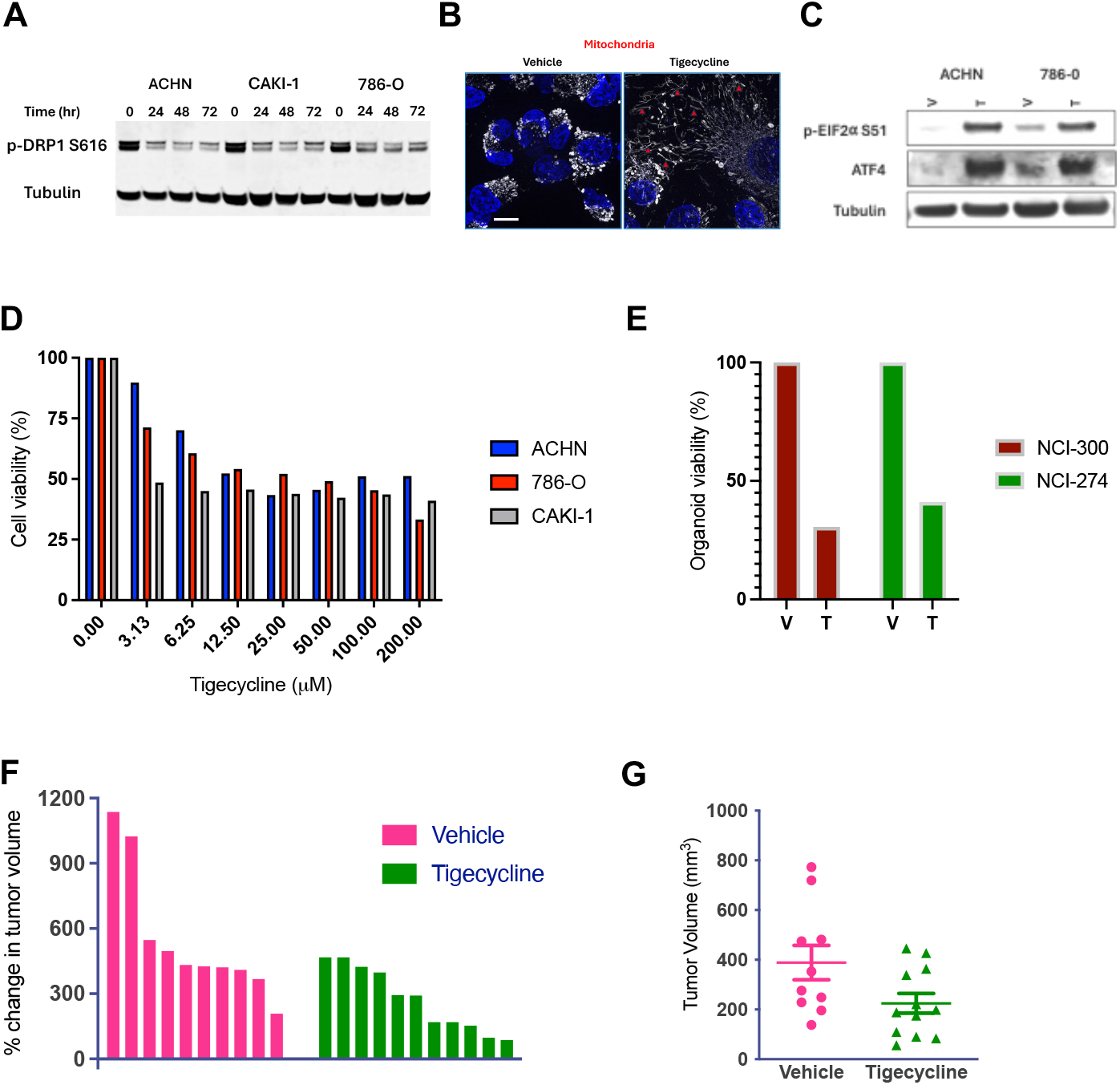
Tigecycline demonstrates potent anti-proliferative activity in RCC cell lines, patient-derived organoids, and xenograft models. **(A)** Western blot analysis showing time-dependent reduction in phosphorylated DRP1 (Ser616) in RCC cell lines treated with tigecycline (25 μM) for the indicated time points (0-72 hours). Tubulin serves as loading control. **(B)** Representative immunofluorescence images showing increased mitochondrial elongation (red arrowheads), characteristic of a fusion-dominant phenotype mitochondrial network in ACHN cells following tigecycline treatment (50 μM, 24 hours; Scale bar = 20 μm). **(C)** Western blot analysis of p-EIF2α and ATF4 expression in ACHN and 786-O cells treated with tigecycline (25 μM) for 48 hours. Tubulin serves as loading control. **(D)** Dose-dependent inhibition of cell viability in ACHN, 786-O, and CAKI-1 RCC cell lines following 72-hour exposure to increasing concentrations of tigecycline. Cell viability was assessed using CellTiter-Glo assay and normalized to DMSO control. Representative data from three independent experiments. **(E)** Response of patient derived organoids (NCI300 and NCI274) to tigecycline (25 μM) treatment for 96 hours. Organoid viability was determined by CellTiter-Glo 3D assay and normalized to DMSO control. **(F)** Waterfall plot showing percent change in tumor volume for individual mice in vehicle-treated (magenta) and tigecycline-treated (green) groups. **(G)** Final tumor volumes at study endpoint in vehicle-treated and tigecycline-treated mice. Each dot represents an individual tumor, and horizontal lines indicate mean ± SEM. *p<0.05 by unpaired two tailed t-test.

### Tigecycline Demonstrates Robust Anticancer Activity In RCC Cell Lines, Patient Derived Organoids and human RCC Xenografts

Next, we treated human RCC cells (ACHN, 786-O, and CAKI) with increasing doses of tigecycline and observed a progressive reduction in cell viability **(Figure 3D)**. The calculated IC_50_ values ranged from approximately 6.25 μM for CAKI cells to 25-50 μM for ACHN and 786-O cells. Next, to assess clinical relevance, we evaluated tigecycline’s effect on patient-derived RCC organoids.

Significantly, patient-derived organoid models (NCI-274 and NCI-300) demonstrated marked sensitivity to tigecycline treatment, with both models exhibiting approximately 60-70% reduction in viability compared to vehicle-treated controls **(Figure 3E)**. The comparable response between established cell lines and patient-derived organoids suggests that tigecycline’s inhibition of mitochondrial translation represents a therapeutically exploitable vulnerability across diverse RCC models, including those directly derived from patient tumors that better recapitulate tumor heterogeneity and microenvironment.

We next examined the efficacy of tigecycline *in vivo* in the human RCC xenograft tumor model. Tigecycline treatment markedly attenuated tumor growth compared to vehicle-treated controls, with treated tumors exhibiting substantially lower percent change in tumor volume throughout the treatment period **(Figure 3F)**. Individual growth trajectories reveal consistent growth inhibition across all tigecycline-treated tumors. End-point analysis confirmed the significant therapeutic benefit of tigecycline, with final tumor volumes in the treatment group (mean ≈ 220 mm.^3^) substantially reduced compared to control tumors (mean ≈ 400 mm3; p: 0.0445, unpaired two tailed t-test; **Figure 3G**). These in vivo findings corroborate our *in vitro* observations.

Accordingly, this consistent activity across diverse models provides compelling evidence that tigecycline represents a viable therapeutic strategy in RCC with potential for clinical translation given its established pharmacokinetic and side effect profiles.

### Compartment-Specific Proteomic Analysis Reveals a Novel Metabolic-Epigenetic Link

The molecular mechanisms underlying tigecycline’s anticancer effects beyond direct mitochondrial inhibition remains unclear. Therefore, to gain deeper insight into tigecycline’s mechanism of action and potentially identify novel vulnerabilities that could be therapeutically exploited, we performed comprehensive subcellular proteomic analyses of tigecycline-treated RCC cells. By separately examining mitochondrial and cytoplasmic fractions, we sought to capture compartment-specific protein changes that might reveal previously uncharacterized consequences of mitochondrial translation inhibition.

Comprehensive mitochondrial proteomics revealed dramatic reorganization of the mitochondrial protein landscape following tigecycline treatment in human RCC cells CAKI-1 and 786-0 [21]. As visualized in the circos plot **(Figure 4A, B)**, tigecycline treatment primarily affected four major biological processes: cytoplasmic translation, protein metabolic processes, organic biosynthetic processes, and peptide biosynthetic processes. The most significantly altered process was cytoplasmic translation, suggesting compensatory upregulation of cytosolic protein synthesis machinery in response to mitochondrial translation inhibition. Quantitative analysis of individual mitochondrial proteins **(Figure 4C, D)** demonstrated consistent downregulation of mitochondrially encoded proteins from the electron transport chain complexes, particularly those within the series of ETC proteins (indicated by the blue bars), and aligns with tigecycline’s established mechanism of action as an inhibitor of mitochondrial translation. Furthermore, the uniform reduction across multiple mitochondrially encoded proteins confirms the comprehensive impact of tigecycline on the entire mitochondrial translation apparatus rather than selective effects on individual proteins. These proteomic findings mechanistically validate our metabolic observations and provide molecular evidence that tigecycline effectively disrupts mitochondrial protein synthesis in RCC cells, leading to coordinated cellular responses that attempt to maintain proteostasis through increased cytoplasmic translation while simultaneously experiencing compromised mitochondrial respiratory function.

**Figure 4.**
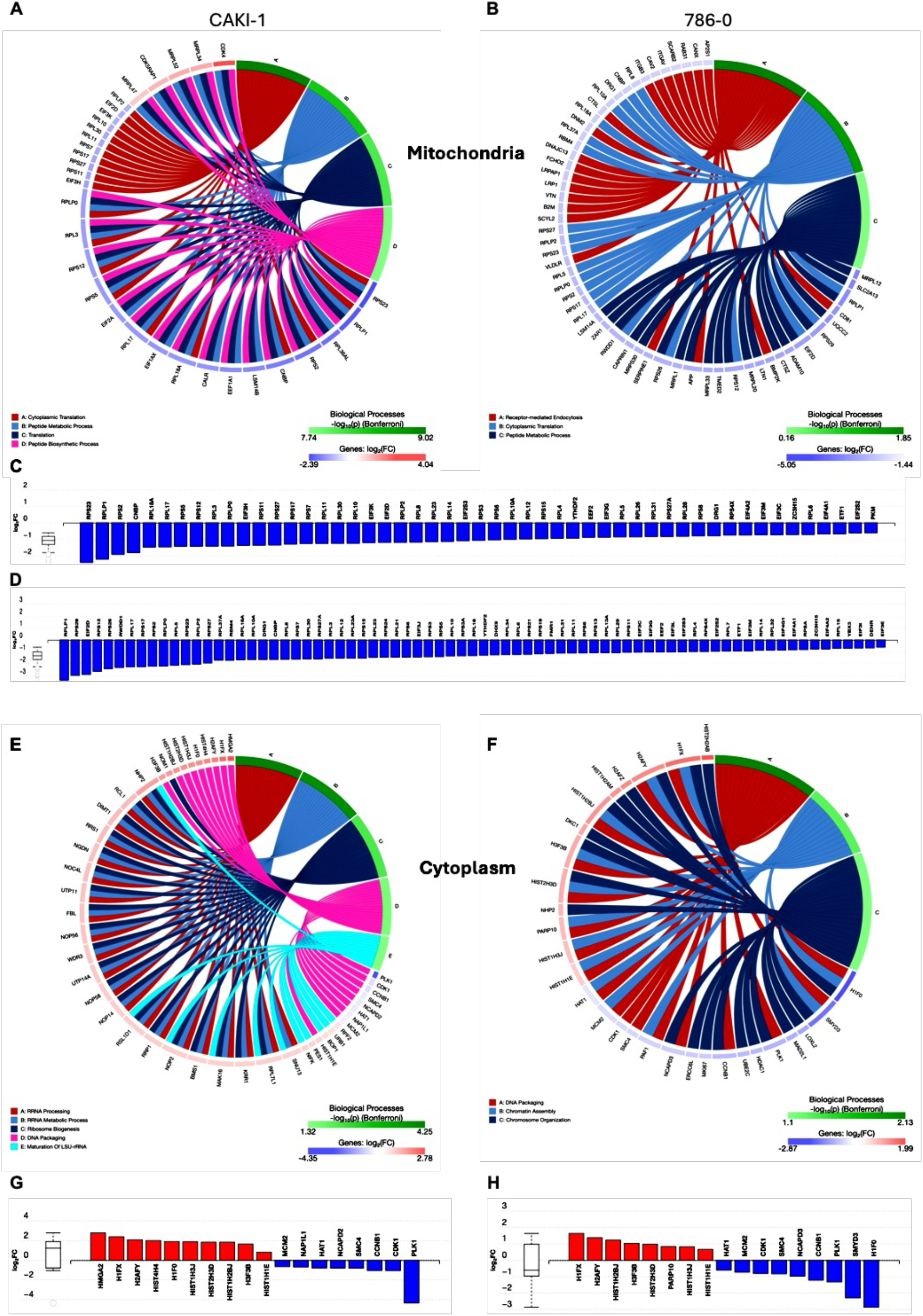
Compartment-specific proteomic analysis reveals distinct molecular signatures in mitochondrial and cytoplasmic fractions following tigecycline treatment. **(A-B)** Circos plots showing enriched biological processes affected by tigecycline treatment in the mitochondrial fraction of CAKI-1 **(A)** and 786-O **(B)** human RCC cells. Colored segments represent major biological processes, with connecting lines indicating protein associations. Color scale (green to red) represents log2 fold change values. **(C-D)** Quantitative analysis of individual mitochondrial proteins in tigecycline-treated CAKI-1 **(C)** and 786-O **(D)** cells compared to DMSO controls. Blue bars represent downregulated proteins (predominantly mitochondrially encoded electron transport chain components). **(E-F)** Circos plots showing enriched biological processes in the cytoplasmic fraction of CAKI-1 **(E)** and 786-O **(F)** cells following tigecycline treatment, revealing distinct patterns compared to mitochondrial fraction. **(G-H)** Quantitative analysis of individual cytoplasmic proteins in tigecycline-treated CAKI-1 **(G)** and 786-O **(H)** cells compared to controls. Red bars represent upregulated proteins (predominantly histone variants), while blue bars represent downregulated proteins (predominantly cell cycle regulators).

### Tigecycline Treatment Drives Dramatic Remodeling of the Cytoplasmic Proteome with histone accumulation and cell cycle perturbation in RCC Cells

Surprisingly, comprehensive cytoplasmic proteomics revealed substantial alterations in protein expression following tigecycline treatment in the same human RCC cells, with patterns distinctly different from those observed in the mitochondrial compartment. As illustrated in the circus plot **(Figure 4E, F)**, tigecycline treatment primarily affected biological processes in the cytoplasm including mRNA processing and rRNA maturation, metabolic processes, nuclear chromatin and chromosome organization. Quantitative analysis of individual proteins demonstrated a pronounced upregulation of multiple histone proteins, including H2AZ, H1FX, H2AFY, H3/H4, H1F0, H3/H4(AU), H3/H3B, H3/H2B, H3/H2B, H2/H1E (red bars, log_2_FC ranging from 0.5 to 2.1). This widespread histone enrichment suggests substantial nuclear chromatin remodeling occurs in response to tigecycline treatment [22,23]. Notably, this histone accumulation occurred concomitantly with significant alterations in cell cycle regulatory proteins **(Figure 4G, H; see also Figure 5G)**. Key regulators of G2/M transition and mitotic progression showed marked downregulation, most prominently PLK1 and cyclin B1 (CCNB1) across multiple cell lines. These proteins orchestrate mitotic entry and progression; consequently, their reduction provides strong evidence for cell cycle arrest at the G2/M phase boundary[24]. Additionally, MCM2 (minichromosome maintenance complex component 2), essential for DNA replication licensing and S-phase progression, was significantly downregulated, further substantiating comprehensive cell cycle disruption. The temporal relationship between histone accumulation and cell cycle perturbation strongly suggests a mechanistic link whereby tigecycline-induced mitochondrial dysfunction triggers cell cycle arrest at a point where histones have already been synthesized for chromosome condensation but cannot be incorporated into chromatin due to blocked mitotic progression [25].

**Figure 5.**
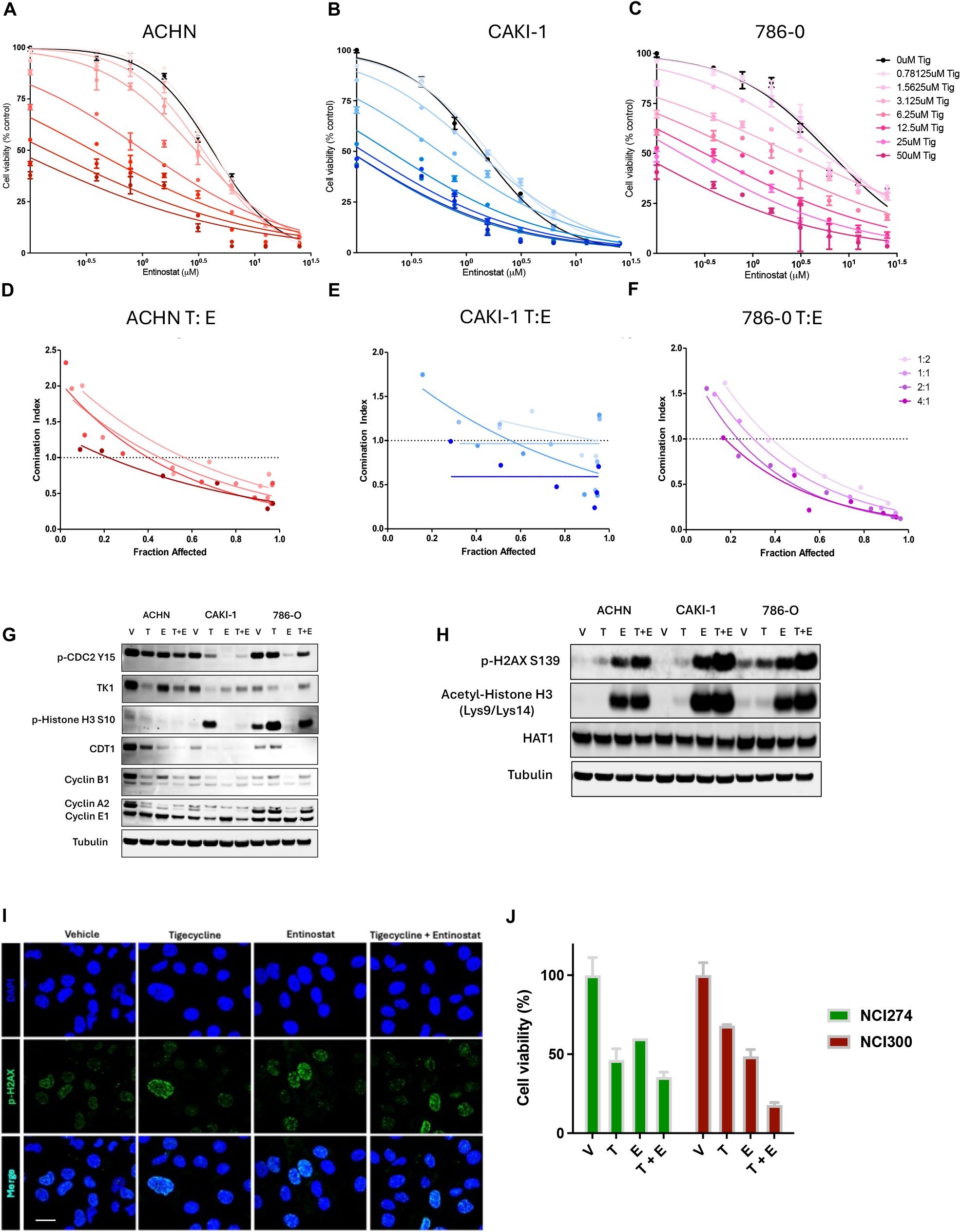
Tigecycline and entinostat exhibit synergistic cytotoxicity in RCC cells through modulation of histone acetylation and cell cycle regulation. **(A-C)** Dose-response curves showing enhanced efficacy of entinostat in the presence of increasing concentrations of tigecycline in ACHN **(A)**, CAKI-1 **(B)**, and 786-O **(C)** human RCC cell lines. Cells were treated with various concentrations of entinostat in the presence of tigecycline (0-50 μM) for 72 hours, and cell viability was assessed using CellTiter-Glo assay. Note the leftward shift of dose-response curves with increasing tigecycline concentrations, indicating enhanced entinostat potency. The best-fit line represents the variable slope (log(inhibitor) versus response). **(D-F)** Combination index (CI) analysis of tigecycline and entinostat combinations at different ratios (1:2, 1:1, 2:1, and 4:1) in ACHN **(D)**, CAKI-1 **(E)**, and 786-O **(F)** cells. CI values below 1.0 indicate synergism, with values approaching 0 representing stronger synergism. **(G-H)** Western blot analysis of cell cycle-related proteins in RCC cells treated with vehicle (V), tigecycline (T), entinostat (E), or the combination (T+E) for 48 hours, with the indicated antibodies. Tubulin serves as loading control.**(I)** Representative immunofluorescence images showing increased γH2AX foci (green) in RCC cells treated with tigecycline, entinostat, or the combination for 24 hours Nuclei were counterstained with DAPI (blue; scale bar = 30 um). **(J)** Viability of patient derived organoids (NCI-274 and NCI-300) treated with vehicle, tigecycline (25 μM), entinostat (2.5 μM), or their combination for 96 hours. Data represent mean ± SEM from three independent experiments.

Taken together, the cytoplasmic proteomic profile stands in marked contrast to the mitochondrial changes, where electron transport chain components were predominantly decreased. The observed histone accumulation represents a previously uncharacterized consequence of mitochondrial translation inhibition in RCC cells and suggests profound nuclear chromatin reorganization occurs as part of the cellular response to mitochondrial dysfunction.

### Combined Tigecycline and Entinostat Treatment Demonstrates Enhanced Efficacy in human RCC cells and Patient Derived RCC Organoids

Based on this mechanistic understanding of histone accumulation and its consequences, we proceeded to test whether targeting this metabolic-epigenetic interface could enhance tigecycline’s therapeutic efficacy. Accordingly, we evaluated whether the histone deacetylase inhibitor entinostat could act synergistically with tigecycline, hypothesizing that HDAC inhibition would further disrupt histone homeostasis in cells already compromised by mitochondrial dysfunction [26] **(Figure 5A-C)**. Our dose-response studies confirmed this prediction, demonstrating that entinostat’s effect on suppressing RCC cell viability were dramatically potentiated in the presence of increasing tigecycline concentrations, resulting in pharmacological synergism characterized by Combination Index values consistently below 1.0 across multiple RCC cell lines **(Figure 5D-F)**. These results demonstrate that mitochondrial translation inhibition via tigecycline creates a cellular state that significantly enhances sensitivity to HDAC inhibition in a dose-dependent manner across different RCC cell lines.

The immunoblot revealed distinct cell cycle perturbations across RCC cell lines treated with tigecycline, entinostat, or the combination **(Figure 5G)**. The decreased phosphorylation of CDC2 at tyrosine 15 observed in tigecycline-treated cells is particularly significant, as this modification serves as a critical regulatory mechanism for mitotic entry. Under normal conditions, CDC2 is maintained in an inactive state through phosphorylation at Tyr15 by Wee1 kinase, and dephosphorylation of this residue by CDC25 phosphatases is required for mitotic progression [27]. The reduced phospho-CDC2(Tyr15) levels following tigecycline treatment indicate attempted activation of the mitotic machinery, but the simultaneous downregulation of cyclin B1 prevents functional CDC2-cyclin B1 complex formation necessary for mitotic progression. The pronounced reduction in thymidine kinase 1 (E2H7Z) expression further confirms inhibition of DNA synthesis machinery, while diminished CDT1 levels indicate suppression of pre-replication complex formation, preventing S-phase re-entry. This coordinated pattern of cell cycle regulator modulation is characteristic of cells arrested at the G2/M transition rather than random dysregulation of cell cycle components. Additionally, the increased phosphorylation of histone H3 at serine 10 in CAKI and 786-O cells following tigecycline treatment. This specific modification is a marker of chromatin condensation during early mitosis and typically peaks during prophase [28]. The elevated phospho-histone H3 Ser10 levels further support our conclusion that tigecycline-treated cells have entered early mitosis but cannot complete cell division, consistent with G2/M phase arrest. The differential regulation of cyclins provides additional mechanistic insight - reduced cyclin A2 (which normally accumulates during S and G2 phases) and diminished cyclin B1 (the principal mitotic cyclin), without compensatory increases in G1-associated cyclin E1, establish a molecular profile consistent with cells that have progressed through S phase but cannot execute mitosis. Importantly, combined tigecycline and entinostat treatment maintained or enhanced these G2/M arrest markers, providing a mechanistic explanation for the synergistic interaction observed between these compounds.

We further observed that the tigecycline and entinostat combination treatment markedly increased phosphorylation of histone γ-H2AX in all three RCC cell lines compared to either agent alone, indicating enhanced DNA damage response[29] **(Figure 5H, I)**. Entinostat’s on-target activity was confirmed by increased acetylation of histone H3 at lysine 9 and 14 residues (AcH3[Lys9/Lys14]), which was further enhanced in combination with tigecycline. HAT1 levels remained unchanged across all treatment conditions. Together, these findings demonstrate that combined inhibition of mitochondrial translation and histone deacetylation synergistically suppresses cell viability in human RCC cells likely through convergent effects on stress response pathways and chromatin organization.

Treatment of patient-derived organoids (PDOs) revealed potent activity of the tigecycline-entinostat combination in clinically relevant models **(Figure 5J)**. In the NCI-300 PDO model single agent tigecycline and single agent entinostat demonstrated moderate efficacy by reducing cell viability. Remarkably, the combination of tigecycline and entinostat exhibited superior anticancer activity compared to either agent alone. This marked enhancement in efficacy suggests synergistic interaction between mitochondrial translation inhibition and HDAC inhibition in patient-derived models that more accurately recapitulate tumor heterogeneity and microenvironment than conventional cell lines. The pronounced effect of the combination treatment on this clinically relevant model provides strong preclinical evidence supporting further investigation of this therapeutic strategy in RCC.

## Discussion

Our study demonstrates that tigecycline effectively inhibits mitochondrial translation in renal cell carcinoma, creating significant bioenergetic deficits and revealing a previously undescribed metabolic-epigenetic interface with therapeutic implications. Tigecycline compromised oxidative phosphorylation capacity and simultaneously impaired glycolytic compensation, creating a profound energetic crisis in cancer cells.

Our compartment-specific proteomic approach provided unprecedented resolution of subcellular adaptations to mitochondrial translation inhibition in RCC cells, revealing distinct molecular signatures that would have remained obscured in whole-cell analyses. The mitochondrial proteome exhibited comprehensive remodeling following tigecycline treatment, with significant downregulation of mitochondrially-encoded electron transport chain components across all complexes, confirming the on-target effects of tigecycline. This mitochondrial disruption triggered compensatory upregulation of cytoplasmic translation machinery, indicating retrograde signaling from mitochondria to nucleus that attempts to maintain cellular homeostasis under conditions of metabolic stress.

In striking contrast, the cytoplasmic proteome revealed an unexpected and pronounced accumulation of multiple histone variants, including H2AZ, H1FX, H2AFY, and various H3/H4 isoforms, concurrent with downregulation of cell cycle regulators including PLK1 and Cyclin B1. This pattern provides mechanistic insight into how mitochondrial dysfunction triggers nuclear events, specifically chromatin reorganization and cell cycle arrest. This metabolic-epigenetic crosstalk reveals a previously undescribed consequence of mitochondrial translation inhibition in cancer cells that bridges traditionally separate domains of cellular biology.

Mechanistically, this histone accumulation likely results from multiple converging processes. G2/M arrest creates a cellular state where newly synthesized histones cannot be incorporated into chromatin during failed mitotic progression [25]. Additionally, mitochondrial impairment alters metabolite availability—particularly acetyl-CoA and α-ketoglutarate—that serve as essential substrates and cofactors for histone-modifying enzymes [30]. The integrated stress response activated by tigecycline further contributes through selective gene expression regulation, potentially favoring histone production while globally attenuating translation. While G2/M cell cycle arrest provides one compelling explanation for this phenomenon, several alternative mechanisms may contribute to this observation. First, mitochondrial dysfunction likely disrupts the availability of key metabolites that serve as substrates and cofactors for histone-modifying enzymes, such as acetyl-CoA pools which are needed for histone acetylation, and is predominantly generated in the mitochondria through pyruvate dehydrogenase and fatty acid oxidation [31]. Second, mitochondrial stress activates retrograde signaling pathways that influence nuclear gene expression programs, potentially including those governing histone synthesis and degradation. The integrated stress response activated by tigecycline selectively upregulates certain genes while globally attenuating translation; this differential regulation might favor histone production over their clearance mechanisms. Third, recent studies have demonstrated that mitochondrial damage can trigger changes in chromatin structure as part of a protective response, potentially involving increased histone synthesis or altered histone positioning [32]. This adaptive mechanism may become dysregulated under sustained mitochondrial dysfunction, leading to histone accumulation.

This metabolic-epigenetic link provides a compelling rationale for our observed synergistic efficacy of combined tigecycline and entinostat treatment. HDAC inhibitors like entinostat target enzymes responsible for removing acetyl groups from histones, promoting a more open chromatin state. By simultaneously disrupting mitochondrial function and histone acetylation status, we exploit a synthetic lethal interaction where cancer cells cannot compensate for both perturbations [33].

Our findings contribute to the growing recognition that metabolic adaptations in cancer create unique vulnerabilities that can be therapeutically exploited. Similarly, VanderHeiden and colleagues have emphasized, cancer cells must balance ATP production with the generation of building blocks for biomass accumulation, creating dependencies that normal cells do not share [34]. The metabolic-epigenetic link we identify expands this concept by demonstrating that targeting cancer metabolism can have profound effects beyond energy production and biosynthesis, extending to fundamental processes like chromatin regulation.

The translational implications of our work are particularly significant. Both tigecycline and entinostat have established pharmacokinetic profiles and safety data in humans, facilitating potential clinical translation of this combination therapy. Especially for patients with advanced RCC who have exhausted standard treatment options, our discovery of this novel metabolic-epigenetic vulnerability offers promising therapeutic potential. Our demonstration that entinostat synergistically enhances tigecycline’s efficacy provides strong validation that this metabolic-epigenetic vulnerability can be therapeutically exploited. HDAC inhibition targets adaptive mechanisms cancer cells employ to manage chromatin organization under metabolic stress. This combination approach overcomes a fundamental limitation of many metabolic-targeting strategies: the remarkable plasticity cancer cells exhibit in nutrient utilization and energetic adaptations. The robustness of this synergism across multiple experimental systems—from cell lines to patient-derived organoids—supports its potential clinical relevance.

Future investigations should focus on characterizing the specific histone modifications altered by mitochondrial dysfunction and identifying biomarkers that predict response to combined mitochondrial and epigenetic targeting. Additionally, exploring how this approach might be extended to other cancer types with mitochondrial dependencies could broadly expand its clinical applicability.

## Author Contributions

Conceptualization, G.V.T; Methodology, K.E.; Software, B.C.; Formal Analysis, K.E., B.C., J.C.,

M. C., M.F., G.V.T.; Investigation, K.E., B.C., J.C., M.C., M.F., G.V.T., Writing – Original Draft Preparation, GVT.; Writing – Review & Editing, GVT, KE, BC.; Project Administration, G.V.T.

## Acknowledgements

We thank Beverly Emerson, Ph.D. for scientific insights

